# Harnessing alkaline-pH regulatable promoters for efficient methanol-free expression of enzymes of industrial interest in *Komagataella phaffii*

**DOI:** 10.1101/2023.12.28.573544

**Authors:** Marcel Albacar, Antonio Casamayor, Joaquín Ariño

**Affiliations:** Institut de Biotecnologia i Biomedicina & Departament de Bioquímica i Biologia Molecular, Universitat Autònoma de Barcelona, Cerdanyola del Vallès, Spain

**Keywords:** heterologous protein expression, methanol-free bioprocesses, phytase production, alkaline stress, promoter mapping, *PHO89*, *HSP12*, *TSA1*, *Komagataella phaffii* (*Pichia pastoris*)

## Abstract

**Background:** The yeast *Komagataella phaffii* has become a very popular host for heterologous protein expression, very often based on the use of the *AOX1* promoter, which becomes activated when cells are grown with methanol as a carbon source. However, the use of methanol in industrial settings is not devoid of problems, and therefore, the search for alternative expression methods has become a priority in the last few years.

**Results:** We recently reported that moderate alkalinization of the medium triggers a fast and wide transcriptional response in *K. phaffii*. Here, we present the utilization of three alkaline pH-responsive promoters (p*TSA1*, p*HSP12* and p*PHO89*) to drive the expression of a secreted phytase enzyme by simply shifting the pH of the medium to 8.0. These promoters offer a wide range of strengths, and the production of phytase could be modulated by adjusting the pH to specific values. The *TSA1* and *PHO89* promoters offered exquisite regulation, with virtually no enzyme production at acidic pH, while limitation of Pi in the medium further potentiated alkaline pH-driven phytase expression from the *PHO89* promoter. An evolved strain based on this promoter was able to produce twice as much phytase as the reference p*AOX1*-based strain. Functional mapping of the *TSA1* and *HSP12* promoters suggests that both contain at least two alkaline pH-sensitive regulatory regions.

**Conclusions:** Our work shows that the use of alkaline pH-regulatable promoters could be a useful alternative to methanol-based expression systems, offering advantages in terms of simplicity, safety and economy.

## INTRODUCTION

Heterologous expression in diverse hosts is an important source for protein production with very diverse uses, from food and beverages to pharmaceuticals. Among the possible hosts, the yeast *Komagataella phaffii* (formerly *Pichia pastoris*) is widely used for heterologous protein production [1,2]. The popularity of this organism derives from several advantageous features, such as high cell density growth in defined media and the ability to perform eukaryotic posttranslational modifications and to release target products extracellularly. In addition, a wide range of genetic modification tools, including genome editing toolkits (CRISPR/Cas9 system), are available for this organism [3].

Expression in *K. phaffii* can be driven from both constitutive and inducible promoters [4]. Examples of the first class are the strong *GAP* and *TEF1* promoters and the somewhat weaker *PGK1*. By far, the most widely employed inducible promoter is that of the *AOX1* gene, which is responsible for most of the alcohol oxidase activity in the cell and allows this organism to grow on methanol as a carbon source. The *AOX1* promoter (p*AOX1*) is induced by methanol and is repressed by carbon sources such as glucose, ethanol, glycerol or acetate [5,6]. The acceptance of this promoter is based on its strength, which allows the production of some secreted recombinant proteins at >15 g/L yield and tight regulation. However, the need to grow on methanol represents a significant drawback because 1) methanol is a hazardous and explosive material that needs special handling, posing a problem in larger-scale industrial processes; 2) it is highly toxic and not suitable for producing certain edible and medical products; 3) its metabolism produces hydrogen peroxide as a byproduct, leading to oxidative stress, which may result in the degradation of recombinant proteins [7–9]; and 4) the use of methanol requires a higher oxygen supply and heat removal than other carbon sources [10,11], thus increasing production costs and making scaling-up difficult. The limitation associated with the use of the *AOX1* promoter has prompted the search for alternative, methanol-free expression systems in *K. phaffii*. Thus, in the last few years, diverse novel regulated promoters that do not require the use of methanol, such as p*THI11* activated by thiamin starvation [12], or the *GTH1* promoter, which is repressed on glycerol and upregulated upon glucose addition [13,14], have been proposed. Other approaches were based on the identification of orthologous promoters from related methylotrophic yeasts [15], or on devising methods for alternative modulation of the p*AOX1* promoter itself [16,17].

Most yeasts, including *K. phaffii*, grow preferentially at acidic pH, and shifting the pH of the medium to moderate alkalinization (pH 7.8-8.2) results in an adaptive response that is mainly mediated by the rewiring of gene expression. Such a response has been characterized in detail in diverse species, such as *Saccharomyces cerevisiae*, *Candida albicans*, and *Aspergillus nidulans* [18–22]. We have recently reported short-term changes in the transcriptional landscape of *K. phaffii* in response to moderate alkalinization of the medium [23]. In this report, we identified several genes that were induced in cells shifted to pH 8.0 at a level comparable to that of strong constitutive genes such as *TEF1* or *GAP1*. We then conceived the idea that such promoters could be an advantageous alternative to p*AOX1*-based expression systems because they would allow the culture of cells on any desired nontoxic carbon source, thus eliminating safety and regulatory issues. Here, we present the use and characterization of three different alkali-regulatable promoters (p*PHO89*, p*HSP12* and p*TSA1*) for the expression of an *E. coli* derived phytase enzyme. This enzyme was selected because, among other interesting characteristics, it represents one of the largest enzyme segments in the feed industry [24]. We show here that strains based on these promoters offer a useful range of strength and, in some cases, could compete with p*AOX1*-based strain used in the industry in terms of potency and fast induction.

## MATERIALS AND METHODS

### Yeast strains and growth conditions

*K. phaffii* X-33 and the new strains generated in this work were grown at 28 °C with continuous shaking (230 rpm/min) in YP medium (10 g/L yeast extract, 20 g/L peptone) supplemented with 20 g/L glycerol (YPGly) or glucose (YPD). YPD plates were supplemented with 200 µg/ml hygromycin B when selection of transformants was needed. The p*AOX1-*based phytase-producing strain (appaT75, CGMCC 12056) was usually grown in YPGly. Low phosphate (low-Pi) medium was prepared by phosphate precipitation from YP as previously described [25]. The estimated Pi concentration in this medium was 0.7-0.8 mM.

### Recombinant DNA techniques

The *Escherichia coli* strain DH5α was used as a plasmid host and grown at 37 °C in LB medium supplemented with hygromycin B (100 µg/ml) when needed for plasmid selection. *E. coli* transformation and standard recombinant DNA techniques were performed as described [26]. Transformation of yeast cells by electroporation was carried out as in [27] with a Gene Pulser electroporator (Bio-Rad).

### Construction of gene reporters and phytase producer strains

The p*TSA1*-GFP reporter used in this work was made as follows. The *TSA1* promoter was amplified by PCR from genomic DNA obtained from the X-33 strain using Q5 polymerase (New England Biolabs) and primers TSA1_BstBI_Fw and TSA1-GFP_Rv_KpnI. The amplified genomic region (-765/+13, from the initiating ATG codon) was digested with BstBI and KpnI and subsequently ligated into the same sites of plasmid pAHYB-GFP [28]. The p*PHO89*-GFP reporter was described in [23].

For the generation of phytase constructs, we used a two-step PCR strategy. First, we amplified the α-factor secretion signal together with the Phytase ORF from the genome of the appaT75 strain. Second, we amplified the *PHO89* (-772/-1), *HSP12* (-749/-1) and *TSA1* (-765/-1) promoters from genomic DNA of the X-33 strain as described above. All these amplicons were designed so that they contained a 3’-region that hybridized with the first 24 nucleotides of the α-factor secretion signal. Finally, each promoter was fused to the phytase ORF by overlapping PCR, and the final product was digested with the restriction enzymes KpnI and NotI. The final products were then cloned at the same sites of the pAHYB-GFP plasmid, replacing the *GFP* reporter. This process was repeated to generate several versions of p*HSP12*- and p*TSA1*-Phytase constructs, with shortened or extended versions of the promoter region used. Constructs were then used to transform *E. coli* cells, and positive clones were selected from LB plates supplemented with hygromycin B. Positive clones were subjected to DNA sequencing of the inserts to verify the absence of unwanted mutations and subsequently linearized by PmeI digestion and introduced into X-33 cells by electroporation to promote integration into the *AOX1* promoter locus. Positive X-33 transformants were selected on YPD plates containing 200 µg/ml hygromycin B. The correct insertion of the constructs was confirmed by PCR. The oligonucleotides used in this work are documented in Additional File 1.

### Posttransformational vector amplification (PTVA)

The PTVA strategy was used to increase the copy number of the p*PHO89*-Phytase construct. To this end, we grew X-33 p*PHO89*-Phytase cells with increasing concentrations of hygromycin B (from 200 to 4000 µg/ml) as described in [28]. However, instead of using agar plates, we adapted the procedure shown in [29], and PTVA was carried out in liquid cultures, increasing the concentration of hygromycin B in the medium every 48 hours.

### GFP reporter assays

For the induction of GFP expression by alkaline stress, cells were grown overnight on YPGly in duplicate, and then cultures were diluted in 20 mL of fresh medium at OD_600_ = 0.5 in baffled flasks (250 ml). Growth was resumed for 16 hours, with the cultures reaching an OD_600_ of approximately 20-25. At this point, cultures were centrifuged for 5 min at 1000 xg, and supernatants were discarded. One set of cultures was then resuspended in the same volume of YPGly supplemented with 50 mM TAPS adjusted to pH 8.0 to induce the expression of GFP. In parallel, the second set was grown in YPGly 50 mM TAPS at pH 5.5 as a non-induced control. Immediately after resuspending the cells, 1 mL samples were taken (t = 0 h) and fixed with formaldehyde as described in [23]. Subsequent samples were taken and fixed at 4, 8, 24, 32 and 48 hours after induction. During cell growth, 1% glycerol (final concentration) was added to the cultures at 8, 24 and 32 hours. In addition, after 24 h of growth, nutrients other than the carbon source were refreshed by the incorporation of 2 mL of 2x concentrated YP medium. The pH of the alkali-stressed cultures was measured after every sampling using a Crison GLP21 pH meter and readjusted to 8.0 (when indicated) by the addition of the appropriate volume of a concentrated KOH solution. The fluorescence of the samples was analyzed in a CytoFLEX (Beckman Coulter) flow cytometer using the FIT-C filter (525/40).

### Phytase expression induction and sample collection

For the induction of phytase expression by alkaline stress, X33 cells carrying p*PHO89*-, p*HSP12*- or p*TSA1*-based phytase constructs were grown overnight on YPGly or YPD as above, and then cultures were diluted in 25 mL of fresh medium at OD_600_ = 0.5 in baffled flasks (250 ml). Growth was resumed for 16 hours, with the cultures reaching an OD_600_ of approximately 20-30. At this point, cultures were centrifuged (5 min, 1000 xg) and resuspended in the same volume of YPD or YPGly supplemented with 50 mM TAPS adjusted to pH 8.0 (unless otherwise indicated) or in noninducing media (see above) as a control.

For p*AOX1*-controlled phytase expression, the appaT75 strain was grown overnight on YPGly medium, and then cultures were diluted in 25 mL of fresh YPGly medium at OD_600_ = 0.5. Cultures were grown and processed as above, except that they were resuspended in fresh YP supplemented with 1% methanol. One ml aliquots were taken and centrifuged (3 min, 13000 xg) immediately after cell resuspension in fresh media (t = 0 h). The supernatants were stored at -20 °C for further analysis. This process was repeated at 8, 24, 48, 72, 96 and 144 hours for most assays. The pH of the alkali-stressed cultures was measured and readjusted to 8.0 (or the indicated pH) as above after each sampling. At these same time points, the carbon source, either glucose, glycerol or methanol (depending on the culture), was newly added to a 1% final concentration. Every 24 h, 2 mL of 2x concentrated YP media was also added to the cultures to refresh the nutrients.

For those experiments involving initial clone screening and those carried out for functional mapping of the *HSP12* and *TSA1* promoter regions, multiple colonies from each transformation were grown overnight on YPGly until saturation. From this point, cultures were diluted in 5 mL of YPGly 50 mM TAPS adjusted to pH 8.0 (plus an aliquot in noninducing media) at OD_600_ = 0.2, and growth was resumed. During these tests, neither nutrients nor carbon sources were refreshed, and the pH of the induced cultures was not readjusted to 8.0. For p*TSA1*- and p*HSP12*-based constructs, cells were grown for 48 and 72 hours, respectively. Then, 1 mL aliquots were centrifuged (3 min, 13000 xg), and supernatants were analyzed for phytase activity.

### Phytase activity assay

Determination of phytase activity was performed essentially as described in the ISO 30024:2009 protocol (https://www.iso.org/standard/45787.html) but scaling down the final volume to 0.4 ml. Briefly, the appropriate dilutions of the fermentation medium were incubated for 30 min at 37 °C in the presence of 5 mM sodium phytate (Sigma‒Aldrich, #68388, inorganic phosphorus <0.1%). The released phosphate was determined by reaction with a mixture of ammonium heptamolybdate and ammonium vanadate reagents plus nitric acid and measuring the A_405_ in a Multiskan Ascent plate reader (Thermo). Phytase activity (U/ml) was calculated using the slope of a previously generated standard curve. One unit is defined as the amount of enzyme that releases 1 µmol of inorganic phosphate from phytate per minute at 37 °C.

### Other techniques

For SDS-polyacrylamide gel electrophoresis analysis, a volume of the phytase-containing supernatants was mixed with 4x SDS‒PAGE loading buffer (200 mM Tris–HCl pH 6.8, 8% sodium dodecyl sulfate (SDS), 0.4% bromophenol blue, 40% glycerol and 20% β-mercaptoethanol) so that the final concentration was 1x. Subsequently, the mixture was heated at 95 °C for 5 min, and 5 µL of the mixture (equivalent to 3.75 µL of medium supernatant) was resolved by SDS‒PAGE at a concentration of 10% acrylamide/bis-acrylamide (37.5:1 ratio). Gels were stained using a Coomassie Blue solution (10% acetic acid, 25% isopropanol and 0.25% Coomassie Brilliant Blue R250, AMRESCO).

Determination of copy number of the heterologous gene was carried out by quantitative PCR (qPCR) analysis as described in [30] using the single-copy gene *ARG4* as reference. Genome size was taken from [31]. qPCR reactions were carried out with the SsoAdvanced™ Universal SYBR^®^ Green Supermix (Bio-Rad) in a Bio-Rad CFX96 apparatus and run by duplicate in two independent amplifications.

Identification of putative transcription factor-binding sites in the promoters of *HSP12* and *TSA1* was performed with the matrix-scan tool from the (RSAT) server [32] using the JASPAR core nonredundant fungi collection [33] and a *p* value threshold of 10^-4^.

Statistical significance was calculated by the unpaired Student’s t test.

## RESULTS

### Long-term expression of GFP from the *PHO89* and *TSA1* promoters in response to alkalinization

The gene *PHO89* (PAS_chr2-1_0235) encodes a high-affinity Na*^+^*/Pi plasma-membrane transporter needed to adapt to situations of phosphate scarcity. Our previous work with the pPHO89-GFP reporter [23] was restricted to rather short time-course experiments (4-6 h after induction by alkalinization). We considered it necessary to test whether the strong induction observed could be maintained or even increased during prolonged cultures. To this end, we initiated the cultures as described in the Materials and Methods section and established two sets of alkali-induced cultures. The first one was induced at time zero, and growth was continued without further modification of the pH (reaching a value of 6.2 at the end of the experiment). In contrast, in the second set, pH was monitored at different times, and pH 8.0 was restored by the addition of potassium hydroxide (Figure 1A). As can be observed, GFP production in the first set increased steadily until 24 h after induction and then decreased gradually. In contrast, maintenance of alkaline stimulation resulted in a continuous increase in the GFP content, reaching a difference of approximately 1.8-fold at the end of the experiment when compared with cells for which stimulation was not sustained over time. We then tested the response of the p*TSA1*-based reporter restoring pH 8.0 as for p*PHO89-GFP*. The *K. phaffii* gene *TSA1* (PAS_chr2-2_0220) encodes a thioredoxin peroxidase known to be involved in oxidative stress responses. As shown in Figure 1B, substantial GFP production was already detected 4 h after induction, and GFP levels increased throughout the entire experiment, reaching values approximately 25% higher than those detected for the *PHO89* promoter. Thus, recursive maintenance of alkaline pH in the medium greatly improved GFP production driven by the tested alkaline pH-sensitive promoters.

**Figure 1.**
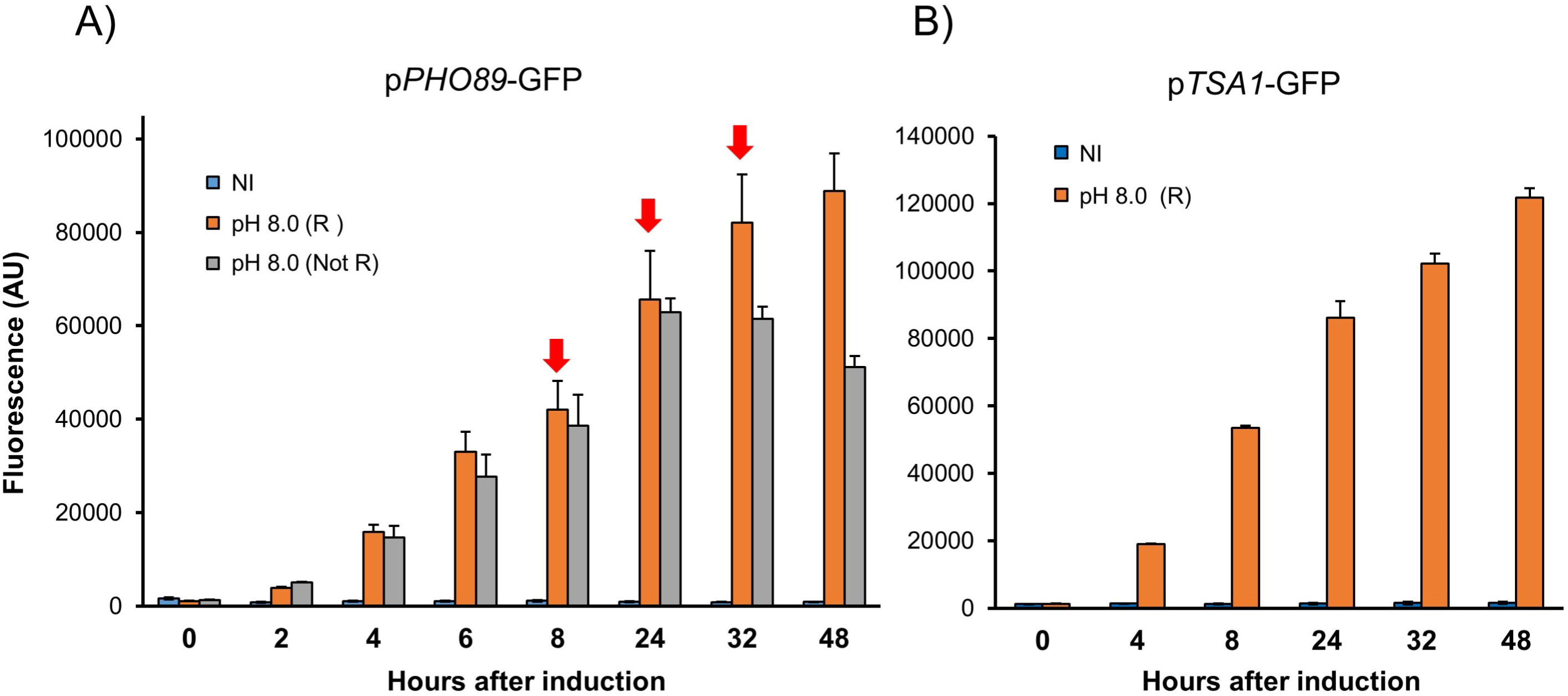
Long-term GFP expression from the *PHO89* and *TSA1* promoters in response to alkalinization. A) A strain with an integrated p*PHO89*-GFP reporter was grown as indicated in the Materials and Methods. At time 0, the cells were resuspended in medium buffered at pH 5.5 (non-induced, NI) or pH 8.0 (induced). The pH of a set of induced cultures was readjusted (R) to pH 8.0 at specific times (denoted by the red arrow), whereas the pH of a parallel set (Not R) was not modified. Samples were fixed, and fluorescence was determined by flow cytometry. Data are presented as the mean ± SEM from 4 to 8 independent cultures. B) A p*TSA1*-GFP reporter strain was cultured as in panel A. In this case, the pH of the induced cultures was readjusted (R) to pH 8.0 after 8, 24 and 32 h of culture. Fluorescence data are the mean ± SEM from 4 independent cultures.

### Expression of phytase from the *TSA1* promoter

The observation that maintenance of the stimulus was beneficial for protein production and that the ratio of detected GFP in induced *vs* non-induced cells with each promoter was very favorable (over 80-fold increase) prompted us to test the ability of these promoters, plus an additional alkaline-pH sensitive promoter (p*HSP12*), to drive efficient expression of the industrially relevant enzyme phytase. In our previous work analyzing the short-term response upon alkalinization, we found that *TSA1* mRNA levels could match those of the powerful constitutive *GAP* promoter, particularly in cells grown on glucose [23]. Therefore, we constructed strains expressing, under the control of the *TSA1* promoter, a phytase form that was engineered to be secreted into the medium and exposed these cells to pH 8.0 using either glycerol or glucose as a carbon source. Twenty transformants were grown in small (5 ml) scale cultures as described in Materials and Methods and tested for production of phytase. Three of the most productive clones were selected and characterized further. Copy number analysis indicated that they carried out two copies of the phytase construct. As shown in Figure 2A (for glycerol) and Supplemental Fig. 1 (for glucose), phytase activity could be detected 8 h after alkalinization for all three clones selected. However, in contrast with what could be expected from the short-term transcriptomic data, the amount of secreted activity was higher (approximately 1.5-fold) for cells grown on glycerol. In any case, the activity produced from the *TSA1* promoter was approximately 25% of that obtained from the reference p*AOX1* strain growing on methanol. We then investigated the sensitivity of the promoter to different levels of stress. To this end, we carried out cultures as in Figure 2A, but using a range of alkalinization from slight (pH 7.4) to moderate (pH 8.2). As shown in Figure 2B, even the lower pH was able to trigger a noticeable production of phytase, albeit the response was somewhat delayed (compare activities at t=8 h). The responses at pH 7.6 and 7.8 were rather similar and somewhat higher than at pH 7.4, and they were further increased at the highest pH values tested. These results suggest that modulation of the stress signal may be used as a tool for the regulation of this promoter.

**Figure 2.**
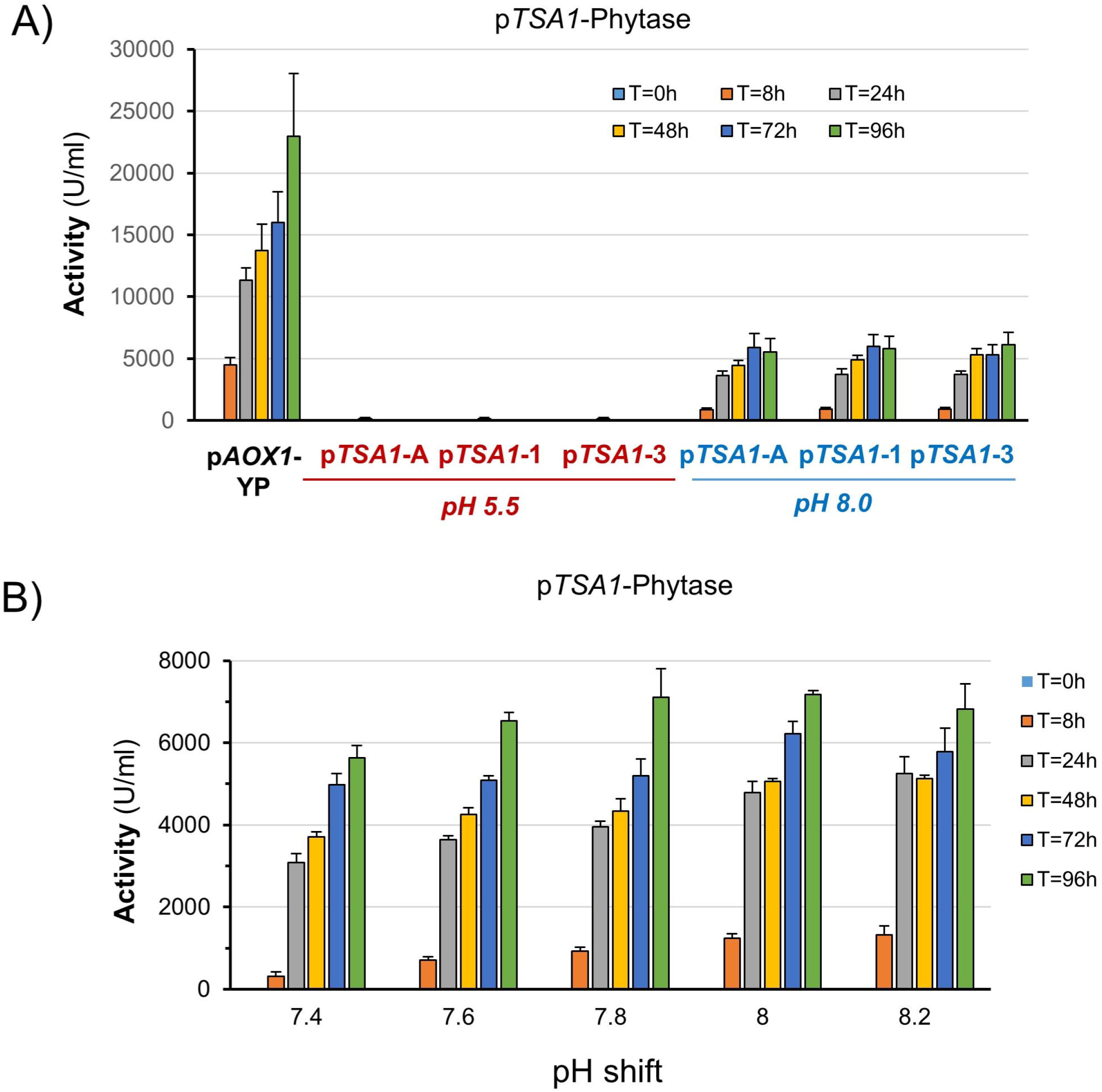
Alkali-induced production of phytase from the *TSA1* promoter. A) Three independent clones containing an integrated p*TSA1*-phytase construct were grown on YP with glycerol as the carbon source and processed as described in the Materials and Methods. Cells were shifted to pH 8.0 at time 0 (or maintained at pH 5.5), and cultures were subjected to periodic restoration of the pH and carbon source. Samples were taken at the indicated times, and phytase activity was determined after appropriate dilution with water. The activity produced by a p*AOX1*-phytase strain grown on YP medium is included for comparison. Data are presented as the mean ± SEM from 3 to 5 independent cultures for each clone. B) The pTSA1-1 clone was processed as in panel A except that cells were shifted at the indicated pH values. Data are the mean ± SEM from 3 independent cultures.

Remarkably, the production of phytase was virtually null in cells grown at acidic pH, suggesting that alkalinization led to positive inputs to the *TSA1* promoter or/and released it from repressive activities. To gain further information, we prepared constructs expressing phytase from different versions of the promoter progressively shortened from the 5’ end (Figure 3A). As shown in Figure 3B, removal of a region spanning from -765 to -459 resulted in a strong decrease in the response to alkaline pH, which was reduced approximately 4-fold, and this behavior was not modified by further deletions of the promoter (to note that the experimental conditions of this experiment are very different from those employed for Figure 2A, leading to a lower phytase production). This result suggested that the functional region relevant for the alkaline pH of the *TSA1* promoter could be restricted to a region of nearly 300 nt spanning from positions -765 to -459. To further refine the analysis, we created four additional deletions (a to d) covering this 300-bp region and generated the corresponding strains. As shown in Figure 3C, the response to alkaline pH for all four strains was reduced to roughly one-half, and at the same time, the phytase activity released by unstimulated cultures became noticeable, suggesting some derepression of the promoter. As a result, the induced/non-induced ratio, which was approximately 200-fold for pTSA1-1, decreased to 2- to 2.5-fold for these four newly constructed strains. Interestingly, additional removal of the region spanning from -527 to -458 (yielding pTSA1-2) not only further decreased the amount of secreted phytase activity in alkali pH-stimulated cells but also abolished the production of phytase in unstimulated cultures. These results suggest that the upstream region of the *TSA1* promoter might contain two independent regulatory components sensitive to alkalinization.

**Figure 3.**
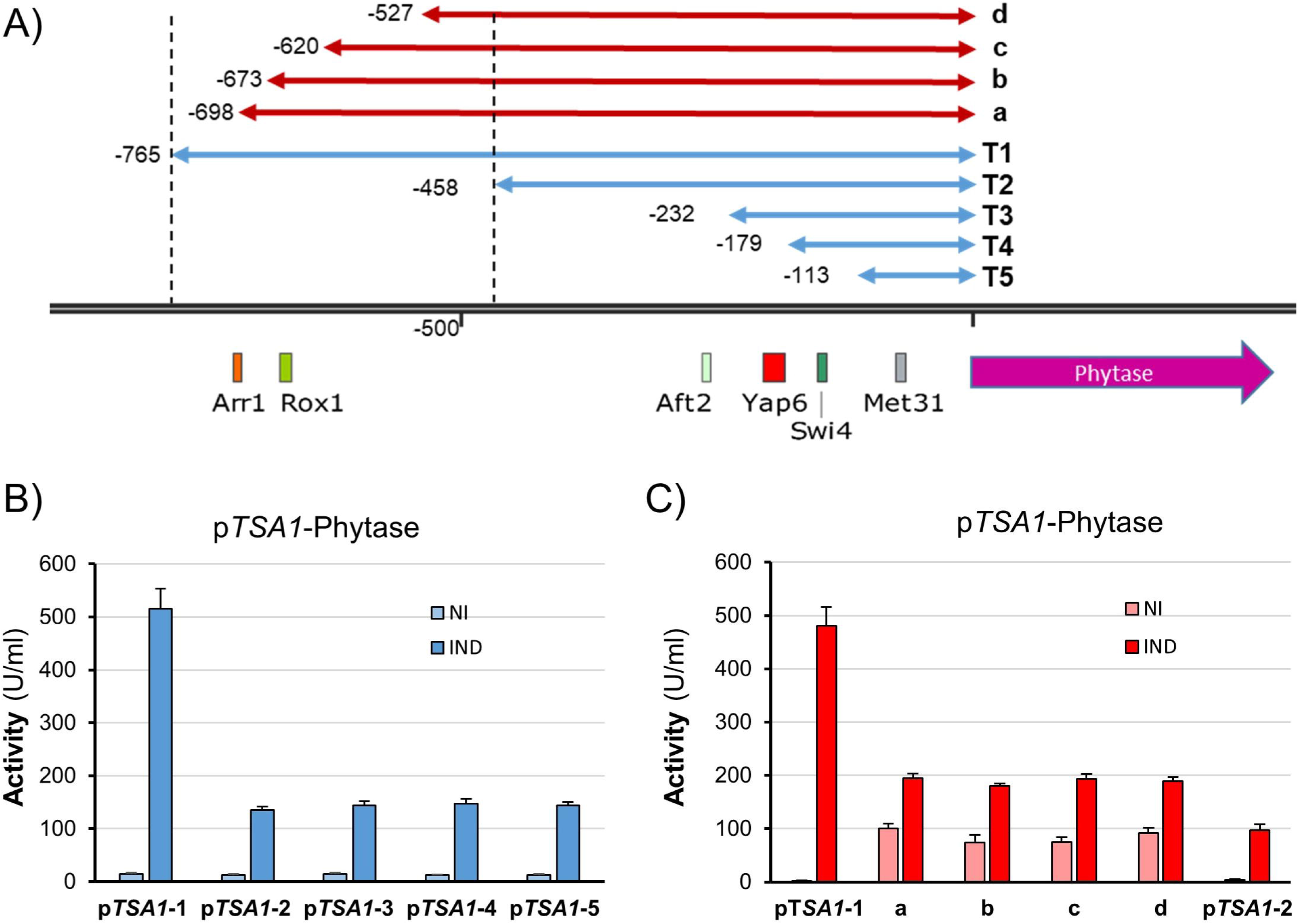
Mapping the pH-dependent activation of the *TSA1* promoter. A new ser of yeast strains that contained the indicated versions of the *TSA1* promoter driving the expression of phytase were created (panel A). The location of predicted transcription factor-binding sites with a *p* value <2x10^-5^ is denoted at the bottom of the cartoon. Cells were grown on YPGly and tested for phytase activity after 48 h of induction (no restoration of pH or nutrients) as indicated in Material and Methods (B and C). Data are presented as the mean ± SEM from 4 (non-induced) or 8 (induced) independent experiments.

### Alkaline induction yields high levels of p*HSP12*-driven phytase activity

*HSP12* (PAS_chr4_0627) encodes a small heat shock protein that is known to respond to diverse stresses. Upon selection of productive clones, as described above, we present in Figure 4A the time course of phytase production for three independent clones expressing phytase from a version of the *HSP12* promoter starting at position -749 relative to the initiating Met (denoted by a red inverted triangle in Figure 4B). The phytase ORF was found as a single copy. Alkalinization resulted in the rapid release of phytase activity, which reached a maximum at the end of the experiment. It is worth noting that, in this case and for some time-points (48 and 72 h), phytase accumulation in the medium was similar to that of the commercial appT75 strain growing on YP plus methanol. At the end of the experiment, the accumulation of phytase from the *HSP12* promoter was approximately 65% of that obtained from the pAOX1-based strain. However, we observed that the uninduced cells also produced detectable phytase activity (see discussion). Consequently, the induced/non-induced ratio was relatively low (3- to 4-fold, depending on the sampling time).

**Figure 4.**
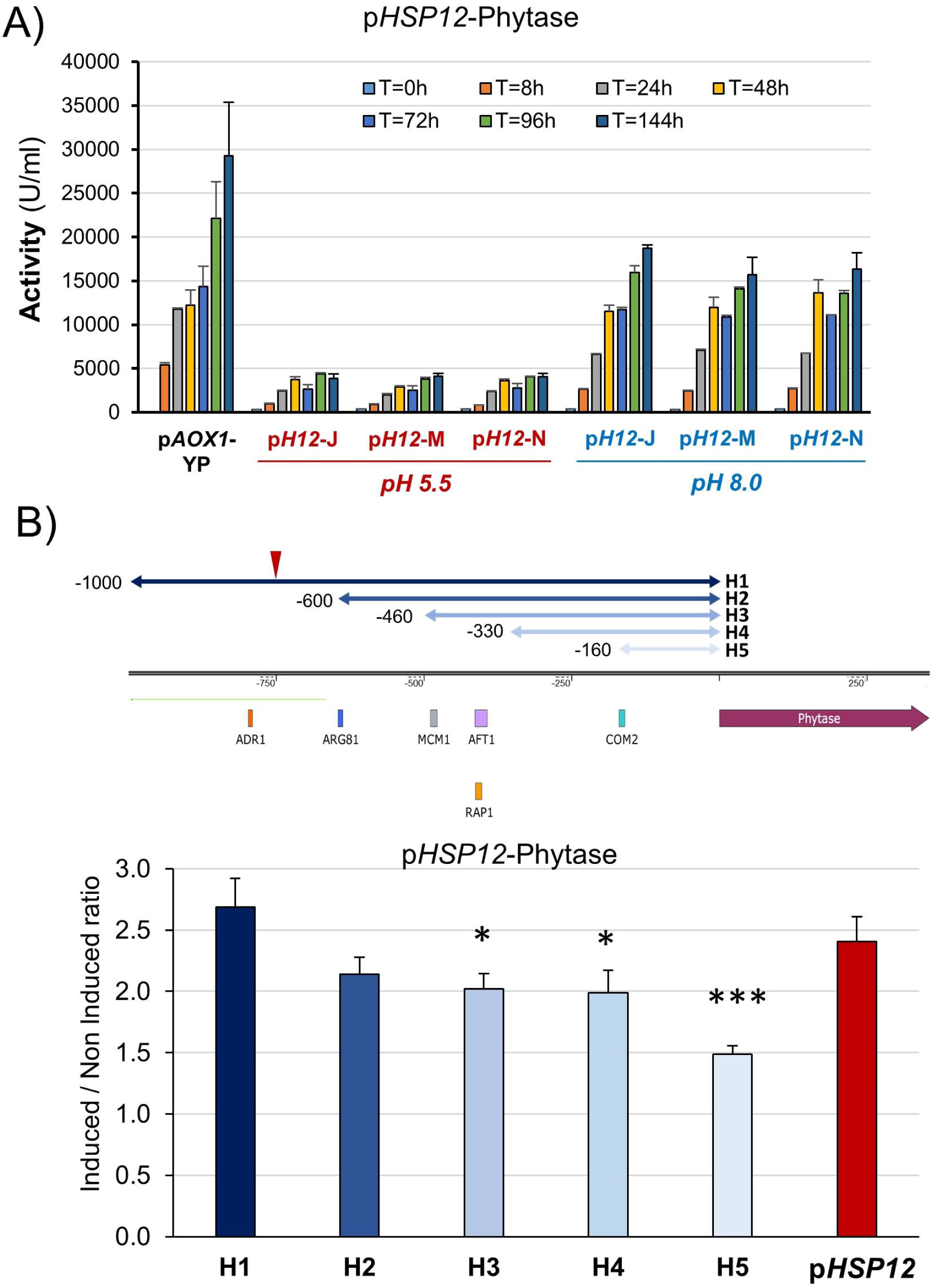
Expression of phytase from the *HSP12* promoter and its functional mapping. A) Three independent clones containing an integrated p*HSP12*-phytase construct were processed as in Figure 2A, and phytase activity was determined in the medium supernatant at different times after induction. Data are the mean ± SEM from 2 or 3 independent cultures for each clone. B) Constructs containing diverse versions of the *HSP12* promoter fused to the phytase ORF were integrated into the genome. The inverted red triangle denotes the position of the 5’ end of the construct assayed in panel A (denoted as pHSP12 in the bar graph). Predicted transcription factor-binding sites (with a *p* value <2x10^-5^) are mapped at the bottom of the cartoon. Phytase activity was determined after 72 h of induction for all strains, and the induced/non-induced ratio was calculated. Data are presented as the mean ± SEM from 8 independent cultures for H1 to H5 and 4 cultures for pHSP12. *, *p*< 0.05, ***, *p*< 0.001.

We wanted to identify possible regions in the *HSP12* promoter that could be sensitive to alkalinization. To this end, we prepared five different versions of the promoter, starting from position -1000 (that is, extending approximately 250 nucleotides upstream in comparison with the promoter present in the strain described above). As shown in Figure 4B, the largest region (H1) provided an induced/non-induced activity ratio of 2.7, quite close to that observed for the p*HSP12*-749 version (note that culture conditions differ from those of Figure 4A). Deletion of the upstream 400 nucleotides (H2) resulted in a noticeable decrease in the induced/non-induced activity ratio, albeit it was not considered statistically significant (*p* value 0.08). Further reduction to positions -460 or -330 (H3 and H4) yielded a similar (and significant) value (≈ 2.0). Finally, when only the upstream 160 nt were included (H5), we observed a further decrease in the activity ratio (1.5). Notably, this last version still provided robust levels of basal activity. From these results, it can be deduced that at least two independent regions sensitive to alkalinization exist in the *HSP12* promoter.

### Effective expression of phytase from the *PHO89* promoter

We prepared a construct in which a promoter region of the same size used for GFP expression was hooked to the phytase ORF and integrated into the yeast genome. As shown in Figure 5A, for three different clones, alkalinization allowed the accumulation of phytase activity to values similar to those attained with the *HSP12* promoter, reaching levels of approximately 50% in comparison with the reference p*AOX1* strain. However, in this case, the activity in the control cultures was virtually undetectable, yielding an induced/non-induced activity ratio of > 1000-fold. Previous work using the GFP reporter suggested that the *PHO89* promoter could be stimulated effectively at pH values lower than 8.0 [23]. Therefore, we carried out a dose-dependence study using one of the clones characterized above (clone P89-J) and tested the production of phytase in a range of pH values from 7.4 to 8.2. As observed from Figure 5B, even the lowest pH tested was able to substantially induce the expression of phytase. Production was increased in cells cultured at pH 7.6 (≈ 1.8-fold at 72 h), although a further rise in the intensity of the stimulus was not reflected in higher enzyme production.

**Figure 5.**
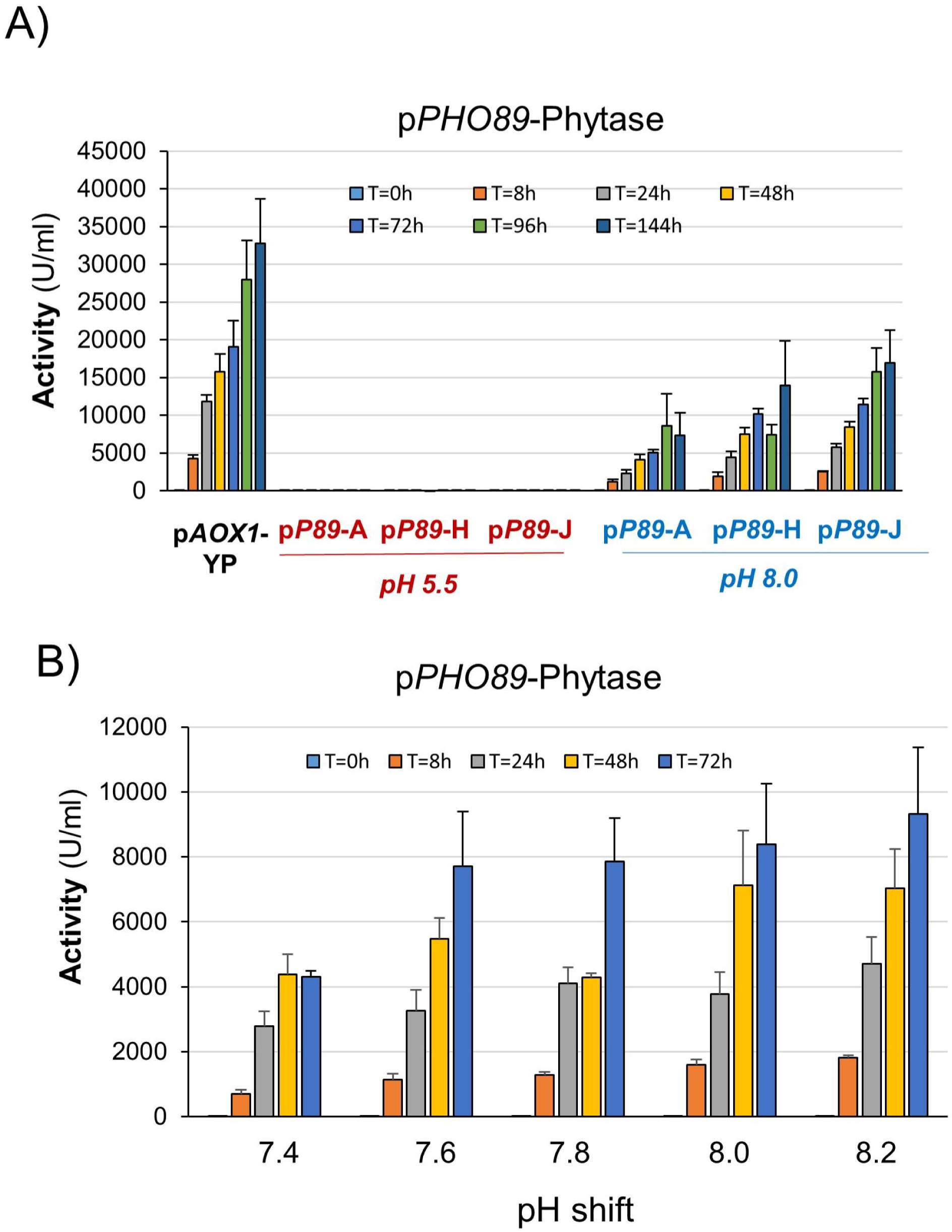
Tightly regulated, alkali-induced expression of phytase from the *PHO89* promoter. Three independent strains, pP89-A, pP89-H, and pP89-J), containing an integrated p*PHO89*-phytase construct, were treated as in Figure 2A, and the produced phytase activity was determined. Data are the mean ± SEM from 3 or 4 independent cultures for each clone. A) Strain pP89-J was induced by resuspending cells in medium buffered at different pH values (from 7.4 to 8.2), which was readjusted (as appropriate) during the experiment. Data are the mean ± SEM from 2 or 3 independent cultures.

The exquisite response of the *PHO89* promoter to alkaline pH prompted us to attempt to improve phytase production by means of the posttransformational vector amplification approach (see Material and Methods). We started with clone P89-J, and at the end of the process, we obtained dozens of clones able to proliferate in the presence of 4000 µg/ml hygromycin (20-fold higher than that used for initial clone selection). After screening for phytase production, we selected three evolved strains (J5, J6 and J7) for additional time-course experiments. As presented in Figure 6A and B, all clones exhibited higher production than the parental P89-J strain (from 1.6 to 3-fold) at all times tested. Such improvement could be explained by the observation that the evolved strains carried 2-3 copies of the phytase-encoding cassette, whereas the p*AOX1*-based strain carried a single copy. The enhanced production of the evolved strain could also be assessed by SDS‒PAGE analysis of the production media (Supplemental Fig. 2), which also shows that phytase is virtually the only protein detected in the supernatant of the cultures. Importantly, the amount of phytase produced was consistently higher for the pH-regulated strains than for the p*AOX1*-based reference strain, with a maximum increase ranging from 2 to 2.5-fold after 72 h of culture (Figure 6B). This difference could be confirmed by SDS‒PAGE analysis. Figure 6C compares the production of phytase from the reference p*AOX1*-based strain and the evolved P89-J6. The amount of protein correlated well with the measured activity. Scanning of the gels and integration of the phytase signals in comparison with known amounts of BSA suggests that the maximal production phytase in the *AOX1*-based strain was 0.59 mg/ml, whereas in the evolved J-6 strains it was 1.32 mg/ml, indicating a specific activity of ̴ 50000 U/mg of protein.

**Figure 6.**
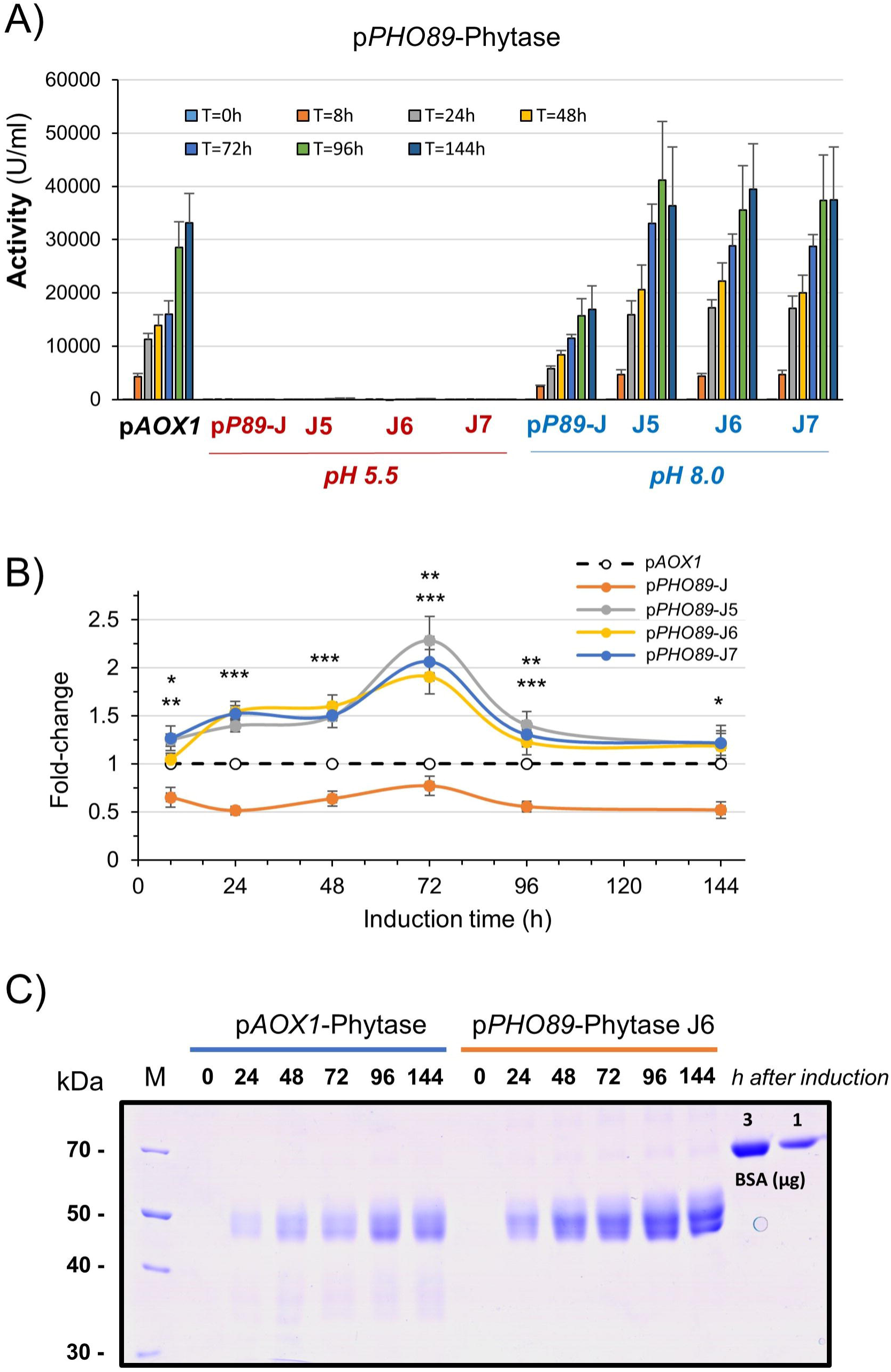
An evolved p*PHO89*-phytase strain surpasses the production levels of the p*AOX1*-phytase strain. A) Three evolved p*PHO89*-phytase strains (J5, J6 and J7), derived from pP89-J, were tested for phytase production as shown in Figure 2A. Data are presented as the mean ± SEM from 3 to 5 independent cultures for each strain. B) The production level of parental pP89-J and the three evolved strains was compared for each time point with that of the p*AOX1*-based strain, taken as the unit. *, *p*< 0.05; **, *p*< 0.01; *** *p*<0.005. C) SDS‒PAGE analysis of the medium recovered from cultures of the p*AOX1*-based and the evolved pP89-J6 strain. Samples corresponding to 3.75 µl of medium were run on 10% polyacrylamide gels and stained with Coomassie Blue. Specific quantities (1 and 3 µg) of bovine serum albumin (BSA) were included in the gel for comparison of the amounts of expressed proteins. M denotes molecular mass standards.

### Phosphate limitation improves alkaline pH-triggered expression from the *PHO89* promoter

*PHO89* encodes a putative Na^+^/Pi high affinity transporter and, as expected, a previous report demonstrated that the *PHO89* promoter positively responds to limitation of Pi availability [34]. We wanted to test if a moderate limitation in Pi could reinforce the response of the *PHO89* promoter to alkalinization. To this end, inorganic phosphate was removed from YP medium by precipitation to decrease its level to about 5-fold (0.7-0.8 mM). The time-course experiment shown in Figure 7, using clone P89-J, demonstrates that partial limitation of Pi in the medium strongly potentiates (nearly 3-fold) the production of phytase, likely by increasing transcription from the promoter. This result is interesting because it illustrates the possibility of modulating alkaline pH response, in a promoter-specific fashion, through judicious manipulation of the culture medium.

**Figure 7.**
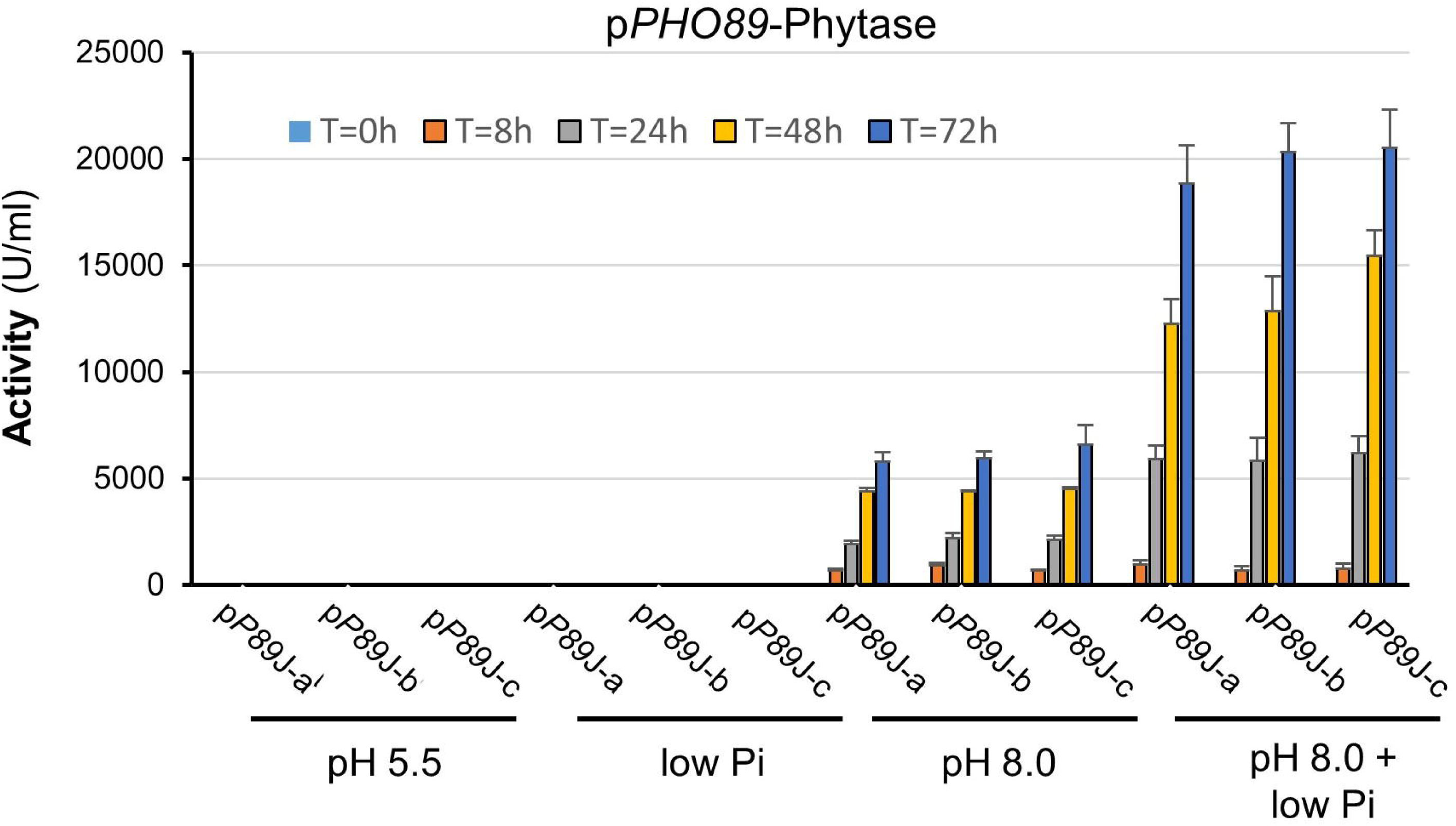
Increased phytase production from the *PHO89* promoter by combination of high pH and phosphate limitation. Three colonies from clone pP89-J (a to c) were grown and tested for phytase production as in Fig. 2A. Low Pi medium was YPD medium partially depleted for Pi (0.7-0.8 mM). Data are presented as the mean ± SEM from three independent experiments for each culture.

## DISCUSSION

We present here three promoters that are plausible, methanol-free alternatives for expression in *K. phaffii* of proteins of industrial interest and that offer a wide range of responses. These promoters were first tested for GFP production and then employed to produce a secreted form of a modified *E. coli* phytase. The choice of phytase as a test enzyme was based on its industrial importance: it is widely used as an additive to monogastric animal feed to improve inorganic phosphate absorption and reduce environmental pollution [35] and its commercial impact since its market value was estimated at USD 517.36 million in 2022 and is growing steadily [36].

Under our experimental setting, the strains based on two of these promoters, p*HSP12* and p*PHO89*, provided expression levels close to or even higher (in the evolved p*PHO89* strains) than those of an p*AOX1*-based strain, while the p*TSA1*-based strain produced phytase at lower levels (approximately a quarter of the yield achieved with p*AOX1* after 96 h). We show that both p*PHO89* and p*TSA1* offer exquisite regulation, with undetectable phytase production when cells grow at acidic pH (Figures 2A and 5B). This might be an invaluable tool when proteins to be expressed could be detrimental to the cell. It is worth noting that in the case of p*TSA1* (and to a lesser extent in *pPHO89)*, it was possible to somewhat modulate the level of phytase expression by adjusting the medium to the appropriate pH (Figures 2B and 5B). This sensitivity to the inducing stimulus provides a way to control the time course of product accumulation. The *PHO89* promoter was proposed earlier as a possible alternative for protein expression, first for the expression of a lipase [34] and later to produce bacterial and fungal phytases [37]. However, in both cases, regulation of the promoter was achieved by using conditions of strong phosphate starvation, which negatively affect growth. In contrast, we did not observe detrimental effects on growth (measured as OD_600_) in cultures readjusted at pH 8.0 at any of the sampling times (Supplemental Fig. 3). Comparison of our results and those provided in reference [37] indicates that, even considering that the enzyme assays are not identical, our production of phytase activity exceeds that obtained by phosphate limitation in those experiments by nearly two orders of magnitude. In the case of p*HSP12*, phytase is also expressed (albeit at lower levels) at acidic pH. This is likely because, in our experimental setting, cells might suffer nutrient limitation for some periods, and it is known that the *HSP12* promoter is induced by glucose starvation in *S. cerevisiae* [38] and is highly and transiently induced upon carbon source limitation in *K. phaffii* [39].

The signaling pathways leading to the regulation of gene expression have been explored in detail in some fungi, such as *S. cerevisiae*, *C. albicans*, *A. nidulans*, and *C. neoformans*. These works have revealed the involvement of several protein kinases [40–46] and transcription factors, such as the conserved PacC/Rim101 [45,47–49], Crz1/CrzA, related to calcium signaling [48,50–54], Pho4, involved in response to phosphate starvation [51,55], Yap1, connected with oxidative stress response [50,56], and likely Mac1 and Aft1, implicated in adaptation to iron/copper scarcity [57,58]. In contrast, virtually nothing is known for *K. phaffii*, except our recent report that the short-term alkali-dependent expression from the *K. phaffii PHO89* promoter was largely dependent on two canonical CDREs, thus strongly suggesting that this response is mediated by the calcineurin/Crz1 pathway [23], as was found earlier for diverse fungi [48,59,60]. In this regard, the response of *PHO89* to alkalinization would be mechanistically independent from that of phosphate limitation, which is likely mediated by putative Pho4-binding sites located at positions -583 and -414 of the *PHO89* promoter [34]. This notion is reinforced by our observation that the combination of alkalinization with slight limitation of Pi, unable by itself to trigger phytase production, strongly potentiates the amount of enzyme recovered from the medium. Such potentiation suggests that the *PHO89* promoter in *K. phaffii*, as we previously demonstrated for the same gene in *S. cerevisiae* [55], integrates two independent signaling pathways. This fact highlights the importance of elucidating the signals elicited by alkalinization and the molecular events occurring downstream, thus leading to the possibility of combination of such responsive elements to create hybrid promoters with enhanced response to alkalinization.

We also show here a functional mapping of p*TSA1* and provide evidence that elements present in two regions, -765/-698 and -527/-458, are important for the alkaline pH response. It is worth noting that none of the stress-responsive transcription factor binding sequences (Rox1, Yap1/6, Aft2) that we could predict to be involved in the alkaline response are located within these regions. The upstream one includes a putative Arr1 binding element (*p* value 1.7E^-5^) at position -723/-716. Arr1 (also known as Yap8) is a transcriptional activator needed for the transcription of genes involved in resistance to arsenic compounds [61]. However, as far as we know, Arr1 has not been previously connected to the response to alkalinization. Our data also show that some responses persist when the promoter is trimmed up to position -113. We identified a Met31 binding site (*p* value < 3.6E^-6^). Met31 is involved in activating genes needed for the incorporation of inorganic sulfate and the synthesis of sulfur-containing amino acids. Remarkably, we recently reported that in *K. phaffii,* alkalinization is characterized by a fast induction of genes involved in sulfate assimilation and Met and Cys biosynthesis [23]. This would support a possible role of Met31 in the alkaline response of the *TSA1* promoter.

The study of the *HSP12* promoter was carried out by generating five different fragments, from position -1000 to -160. We did not trim the promoter further because a recent report showed that a region starting at nt -97 had no activity at all [39]. Our data suggest that this promoter contains two regions sensitive to alkalinization. The first one would lie between nt -1000 and -460 (or possibly -600). This region contains an Adr1-binding sequence (p<1.5E^-5^), which could be recognized by Mxr1, a proposed homolog in *K. phaffii* for yeast Adr1 [62,63].

On the other hand, our results discard the relevance of Rap1, a multifunctional and conserved transcription factor related to oxidative stress conditions in humans and yeast [64,65] that has been identified as important for the expression of catalase (*CAT1*) in *K. phaffii* [66] or Aft1, which in *K. phaffii* has been related to carbohydrate and not to iron metabolism [67]. In contrast, the region between -330 and -160 appears relevant for alkaline induction. This segment contains a putative site for the Com2 transcription factor (*p* value 3.5E^-6^). This factor has been found in *S. cerevisiae* to be necessary for the expression of genes involved in sulfur assimilation (also observed in *K. phaffii* upon alkalinization, see above) when cells are exposed to sulfur dioxide [68]. It must be noted, however, that the sequence recognized by this factor includes a canonical stress-responsive element (STRE, 5’-CCCCT-3’), which is recognized by Msn2/Msn4, and that there is recent evidence for the involvement of these factors in the regulation of *HSP12* in *S. cerevisiae* [69]. In any case, it would be most desirable to investigate the specific signaling pathways governing the transcriptional response to alkalinization in *K. phaffii*.

The use of alkaline pH-regulatable promoters would allow, in principle, the utilization of any carbon source for cultivation of yeast cells. This does not imply that the carbon source would be irrelevant since it might affect the potency of the response, as we show here for the case of the *TSA1* promoter. It is evident that the specific growth conditions for optimal production must be set for each specific promoter and produced protein. In any case, these promoters avoid the use of methanol and hence circumvent the associated problems, such as flammability, toxicity (either by ingestion, inhalation, or absorption through the skin), as well as the requirement for high oxygen supply and concomitant heat production. In contrast, in the expression systems presented here, induction of the desired protein can be achieved without any modification of the medium, simply by the addition of the appropriate amounts of KOH, possibly one of the cheapest chemicals. Potassium hydroxide would be preferable to sodium hydroxide to avoid the accumulation of sodium cations, which are more toxic than potassium [70], in the culture medium. In addition, pH is a very easy parameter to follow in a bioreactor. On-line monitoring can be coupled to the addition of KOH to counteract possible acidification caused by fermentation, thus maintaining the appropriate pH profile for the desired period. Finally, while alkaline-pH regulatable promoters could be used, in principle, to express any protein (preliminary work in our laboratory suggests that it is also suitable for the production of the CRL1 lipase from *Candida rugosa*), the induction conditions would be particularly suitable for secretion of proteins that are unstable at the acidic pH found in standard yeast fermentations, such as methionine adenosyltransferase [71] or the fungal Fsl2 lipase [72]. In any case, the actual competitiveness of the expression systems presented here needs to be evaluated at the bioreactor level and in a pre-industrial setting.

## ADDITIONAL MATERIAL

Additional file 1 (xlsx) Oligonucleotides used in this work.

Additional file 2 (pdf). Supplemental figures 1 (alkali pH-regulated expression of phytase from the *TSA1* promoter in cells growing on glucose), 2 (SDS‒PAGE analysis of the medium recovered from cultures of the parental pP89-J and the evolved pP89-J-6 strains), and 3 (Total cell accumulation, determined as OD_600_, in cultures grown at pH 5.5 and those stimulated at pH 8.0).

## DECLARATIONS

### Ethics approval and consent to participate

Not applicable.

### Consent for publication

Not applicable.

### Availability of data and materials

Any data not presented in the paper are available from the corresponding author on reasonable request.

### Authors’ contributions

MA designed and constructed plasmids and yeast strains and carried out yeast cultures and treatments and protein gels. AC contributed to the design and preparation of DNA constructs, yeast strains and cultures for induction experiments and sampling. JA performed the phytase assays, supervised the work and drafted the manuscript. All the authors have read and approved the final manuscript.

## Acknowledgments

The skillful technical assistance of Montserrat Robledo is acknowledged. We thank Dr. Patricia Puig (Andrés Pintaluba S.A) for advice in the determination of phytase activity.

