## Additional File 2 (supp. Figs. 1-3) for "Harnessing alkaline-pH regulatable promoters for efficient methanol-free expression of enzymes of industrial interest in *Komagataella phaffii*"

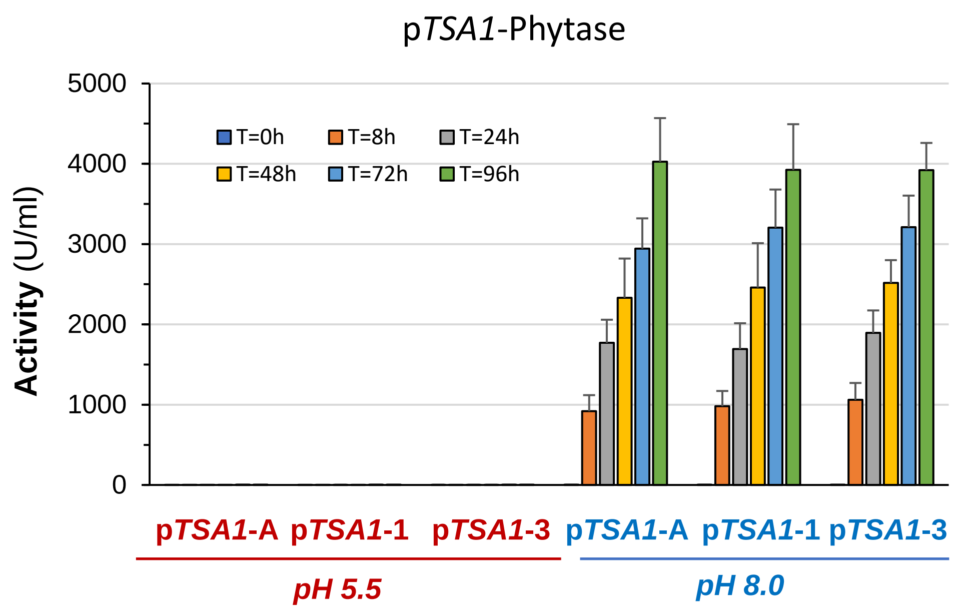


**Supp. Figure 1**. Alkali pH-regulated expression of phytase from the *TSA1* promoter in cells growing on glucose. Three independent clones containing an integrated p*TSA1*-Phytase construct were grown YP with glucose as carbon source and processed as described in Material and Methods. Cells were shifted to pH 8.0 at time 0 (or maintained at pH 5.5), and cultures subjected to periodic restoration of the pH and carbon source. Samples were taken at the indicated times and phytase activity determined after appropriate dilution with water. Data are mean ± SEM from 3 or 4 independent cultures.

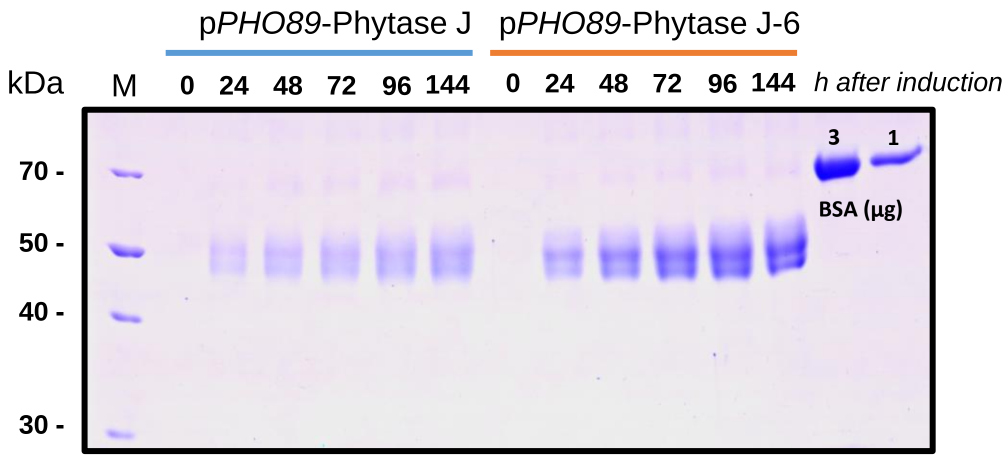


**Supp. Figure 2.** SDS-PAGE analysis of the medium recovered from cultures of the parental p*PHO89*-J and the evolved p*PHO89*-J-6 strains. Samples corresponding to 3.75 µl of medium were run on 10% polyacrylamide gels and stained with Coomassie Blue. Specific amounts (1 and 3 µg) of bovine serum albumin (BSA) were included in the gel for mass comparison. M denotes molecular mass standards.

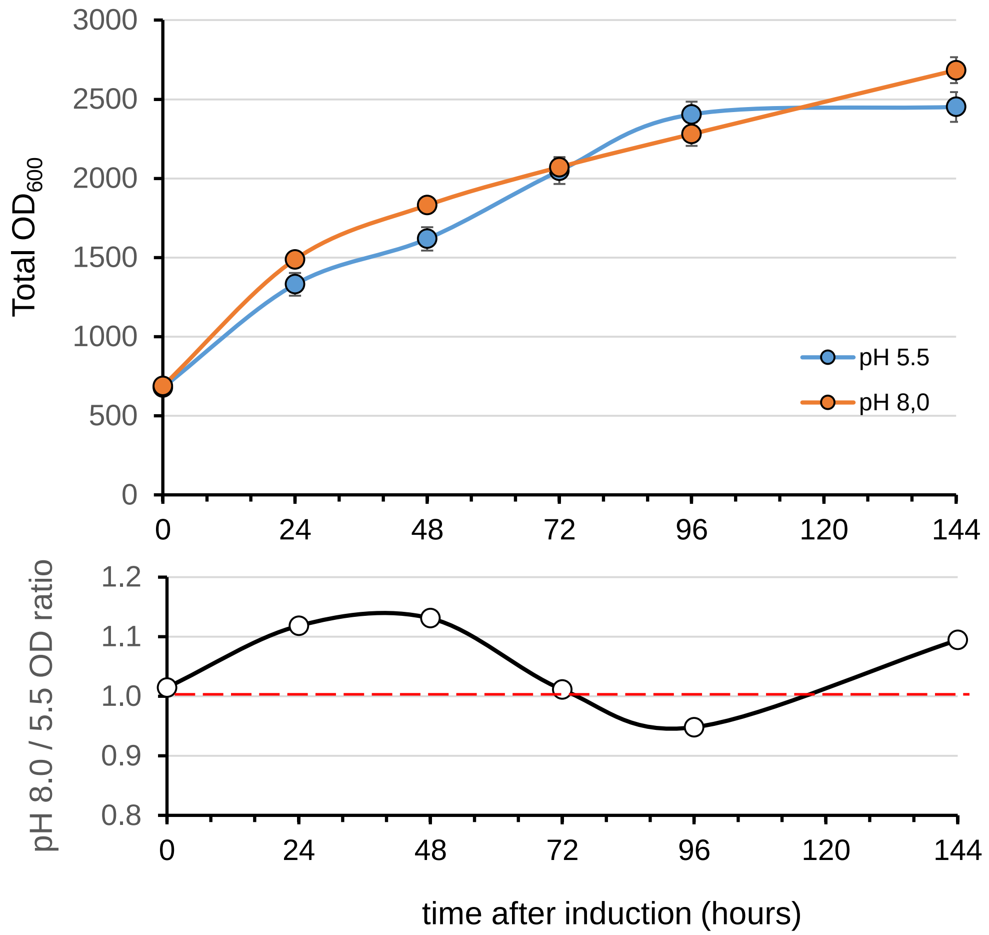


**Supplemental Figure 3**. Upper panel) Total cell accumulation, expressed as $OD_{600}$ x culture volume (ml) in cultures grown at pH 5.5 or regularly shifted at pH 8.0 as described in Material and Methods. Note that along the experiment the sampling procedure results slight differences in the volume in both kind of cultures, so the typical illustration of $OD_{600}$ would not represent accurately the amount of biomass. Data are mean ± SEM from 14 to 26 (pH 5.5) or 26 to 44 determinations (pH 8.0). Lower panel) Calculated ratio of biomass (pH 8.0 / 5.5 ) for each time-point.
